# Molecular and bacterial contamination of a small freshwater effluent used for drinking water

**DOI:** 10.1101/2023.02.28.530486

**Authors:** Carolina Oliveira de Santana, Pieter Spealman, David Gresham, M. Elias Dueker, Gabriel G. Perron

## Abstract

Sewage contamination of freshwater occurs in the form of raw waste or as effluent (at varying levels of treatment) from wastewater treatment plants. Global management of this contamination has focused on detection of live sewage-indicating bacteria in freshwater, drinking water, and irrigation systems. While raw waste (animal and human) and underfunctioning WWTP’s can introduce live enteric bacteria to freshwater systems, most WWTP’s, even when operating correctly, do not remove bacterial genetic material from treated waste, resulting in the addition of concentrated enteric bacterial DNA (molecular contamination), including antibiotic resistance genes, into water columns and sediment of freshwater systems. In freshwater systems with both raw and treated waste inputs, then, there will be increased interaction between live sewage-associated bacteria (untreated sewage) and molecular contamination (from both untreated and treated wastewater effluent), with the potential of increasing antibiotic resistance in the live bacterial populations. To evaluate this understudied interaction between molecular and bacterial contamination in the freshwater environment, we conducted a three-month field-based study of sewage-associated bacteria and genetic material in water and sediment in a freshwater tributary of the Hudson River (NY, USA) that supplies drinking water and receives treated and untreated wastewater discharges from several municipalities. Using both molecular and culture-based bacterial analyses, we demonstrate both treated and untreated sewage influences on water and sediment bacterial communities in this tributary, and water-sediment exchanges of enteric bacteria and associated genetic material with rain events. Furthermore, treated sewage effluent on this waterway serves as a concentrated source of *int1* (antibiotic resistance) genes, which appear to collect in the sediments below the outfall along with fecal indicating bacteria, serving as a possible genetic exchange substrate and a source for future molecular and bacterial water contamination.

**Graphical abstract:** 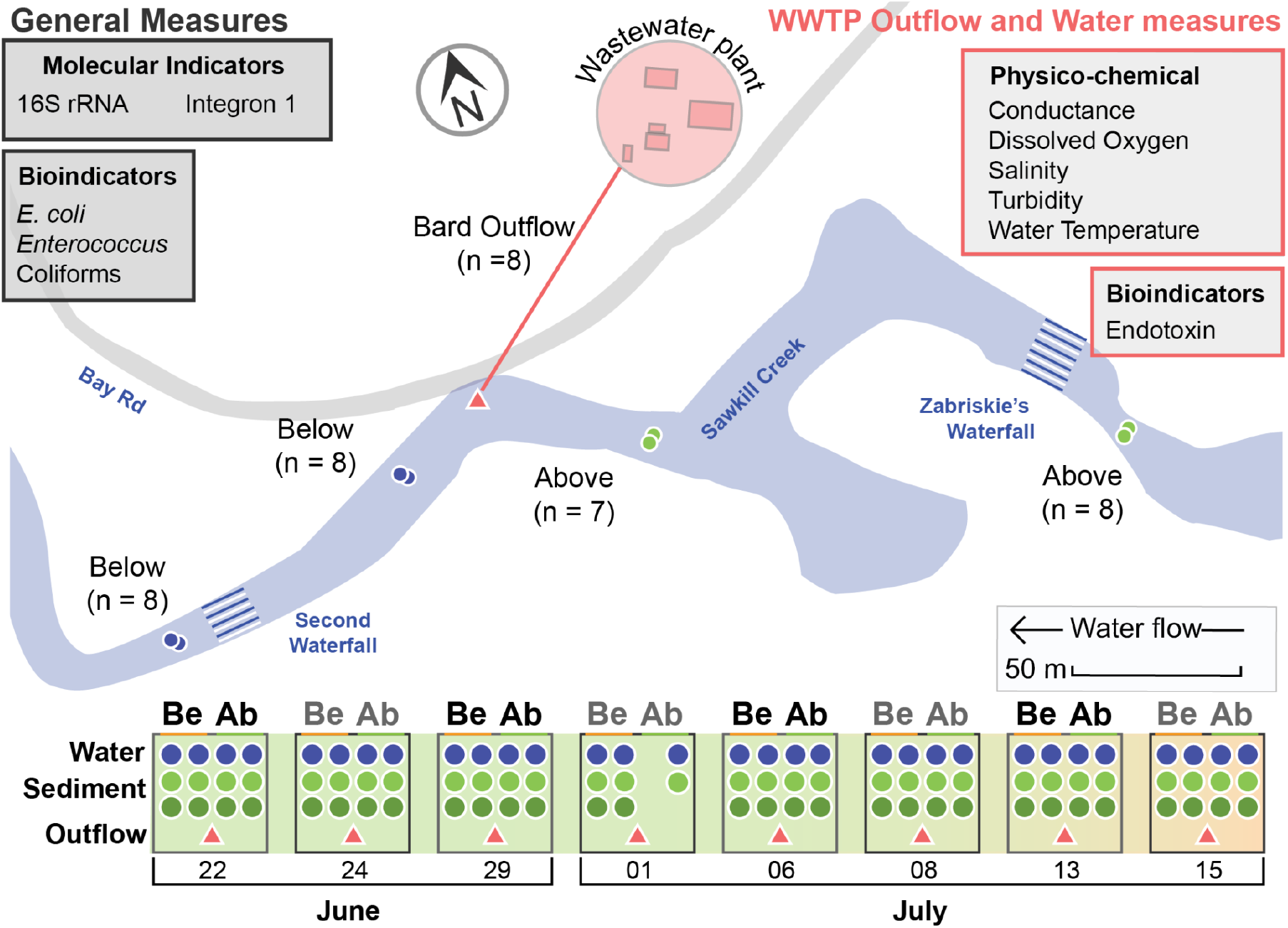

**Highlights:**

- In a model freshwater system used both as drinking water and wastewater disposal,
- Bacterial and genetic material differ between water and sediment compartments
- Live bacteria

## 1. Introduction

Human access to clean water has become one of the most pressing challenges of the 21st century, as climate change accelerates and populations grow without sufficiently adaptable infrastructure (REF). This reality requires careful management of existing freshwater resources and the multiple uses of those resources, which can often conflict. Freshwater streams and rivers around the world, for example, are commonly used for withdrawal of drinking water, bathing and cleaning, and for the disposal of treated and untreated wastewater. The increasing scarcity of freshwater, particularly under extreme weather conditions, only exacerbated the challenges associated with waterborne diseases that can be transferred to humans, which remains a leading cause of death and illness globally (Landrigan et al., 2018).

Public health concerns regarding the treatment of waterborne diseases are further complicated by the spread of antibiotic-resistant bacteria in freshwater environment. Indeed, the prevalence of antibiotic-resistant bacteria in the environment increased globally under the influence of domestic, agricultural, industrial pollution (REFs). Furthermore, polluted environments like freshwater favor the horizontal exchange of antibiotic resistance genes between pathogenic and non-pathogenic bacteria via various mechanisms such as plasmids, phage transduction, and integrons (10.1371/journal.pone.0017038, 10.3389/fmicb.2014.00648, 10.3389/fmicb.2016.00173, 10.3390/w14060938). For these reasons, polluted freshwater environments are now believed to be an important reservoir of antibiotic-resistance genes and a possible site for the emergence of novel antibiotic-resistance bacterial lineages of clinical importance (10.1038/s41579-021-00649-x, <u>10.1099/mgen.0.000455</u>).

To prevent waterborne disease in communities utilizing mixed-use waters, resource managers rely on fecal indicating bacteria (FIB) to detect the presence of untreated sewage in freshwater systems and drinking water systems. While these methods are effective in detecting *untreated* sewage presence in freshwater, they do not detect the presence of *treated* sewage in freshwater, which, among other chemical contaminants, can include concentrated pharmaceuticals {Wilkinson, 2022 #2692}{Cantwell, 2018 #2382}{Rosi-Marshall, 2015 #2693}, concentrated cleaning and personal care products (Montes-Grajales et al., 2017), concentrated endotoxins{Huang, 2011 #2104}, and intact genetic material such as antibiotic resistance genes{Czekalski, 2012 #2694}{Liu, 2018 #2695}.

Furthermore, the intermittent challenge of managing sewage treatment under high-volume events including rainstorms and other extreme weather phenomena results in predictable under-treatment of raw sewage. For example, in New York City, USA, approximately 470 billion gallons of treated sewage are released every year (NYC DEP 2022) into the Hudson River, along with over 20 billion gallons of untreated sewage through combined sewer overflows (CSO) (DePalma, 2007). Extreme weather events present even starker examples of this mixture of treated and untreated sewage in public waterways, as demonstrated by the release of over 10 billion gallons of raw and partially-treated sewage into the Hudson River during Hurricane Sandy in 2012 (Kenward et al., 2013).

The addition of raw sewage to a freshwater system already receiving treated wastewater effluent regularly creates a context for interactions between live sewage-associated pathogens and reservoirs of genetic material, including antibiotic resistance genes, in the environment. Indeed, microbial DNA and bacterial antibiotic resistance genes, including the *intI1* integron cassette, have been detected in both raw sewage and treated effluent in the past (10.1007/s00248-016-0843-4, 10.1016/j.watres.2021.117720, 10.1128/AEM.02766-17) and have even been suggested as treated and untreated sewage indicators (10.1038/ismej.2014.226, 10.1016/j.envint.2019.105372). Crucially, freshwaters, like most other environments, are not monitored for these non-culturable indicators of sewage-associated pollution (10.3390/microorganisms7060180, 10.1021/acs.est.1c08918, Hamilton et al (2022, in press)), but managing their presence is important for the protection of drinking water resources, since genetic material and chemical contaminants are often not removed by most drinking water facilities worldwide. Recreational use of freshwater with high concentrations of treated sewage effluent also potentially exposes users to these contaminants through skin, digestive system, and respiratory contact (REF).

While the extra-enteric ecology of indicator organisms such as Enterococcus, Coliforms, and *Escherichia coli* remains an understudied realm {O’Mullan, 2017 #2152}, strong evidence suggests that these organisms and associated pathogens can and do persist in both the water column and sediments of aquatic systems{Brooks, 2015 #2696}{Brooks, 2021 #2697}{Characklis, 2005 #2698}{Mueller-Spitz, 2010 #2699}. In fact, sediments display impacts of untreated sewage releases much longer than the water column in flowing systems{Mallin, 2007 #2700}, and appear to harbor sewage indicators well past known waste releases{Brooks, 2015 #2696}.

Sediment and water column microbial habitats also are prone to mixing through wind and rain events that disturb and sometimes scour out loose sediments, suspending these sediments in the water column to be moved downstream {O’Mullan, 2017 #2560}. This dynamic sets up the potential for bacteria residing in sediments to interact with bacteria and genetic elements present in the water column, and vice versa.

WWTP effluent releases into freshwater streams impact microbial communities through sand and grit deposition{Wakelin, 2008 #2576}, nutrient enrichment and eutrophication{Howarth, 2008 #2701}, and chemical alterations of water column and sediment pore waters. Sediments below outfalls appear to harbor sewage indicator organisms, suggesting that associated treated sewage elements such as pathogens, genetic material, and pharmaceuticals might also accumulate in sediments near the outflow. These sediments might then serve as a reservoir for bacteria and genetic elements related to pathogenicity and antibiotic resistance. With intermittent disturbance of these sediments, microbial communities that have grown with this mixture would then be distributed further downstream, shifting water column bacterial community structure, seeding new sediments, and generally broadening the influence of the single effluent pipe. Furthermore, in freshwater systems with both raw and treated waste inputs, there may be an understudied potential for interaction between live sewage-associated bacteria (untreated sewage) and sewage-derived genetic material (treated wastewater).

To evaluate this potential, we conducted a three month field-based study of sewage-associated bacteria and genetic material in water and sediment in a freshwater tributary of the Hudson River (NY, USA), (de Santana *et al*. 2022). Like many freshwater systems in the US, this waterway both provides drinking water and receives treated and untreated wastewater discharges from several municipalities. Here, we investigate the possible effects of a small-scale private wastewater treatment plant operated by Bard College on the population structure of bacterial communities found in water and sediment samples collected over eight weeks in the summer of 2015.

## 2. Materials and methods

### 2.1 Study Site

As outlined previously in De Santana et al. (De Santana 2022), this study was conducted on the Saw Kill, a 23.0 km tributary of the Hudson River located at river mile 98 (Fig. 1). The Saw Kill Watershed drains a 57 km^2^ area, which hosts both agricultural and suburban developed land uses (Wikipedia Saw Kill 2022). There are 2 permitted sewage outfalls (NYSPDES permit #’s NY0271420, NY0031925), and one permitted drinking water facility (130,000 gallons per day) along this waterway. Additionally, just downriver from where the Saw Kill pours into South Tivoli Bays (and then to the Hudson River), 7 municipalities pull water from the river proper for municipal drinking water use.

**Figure 1.**
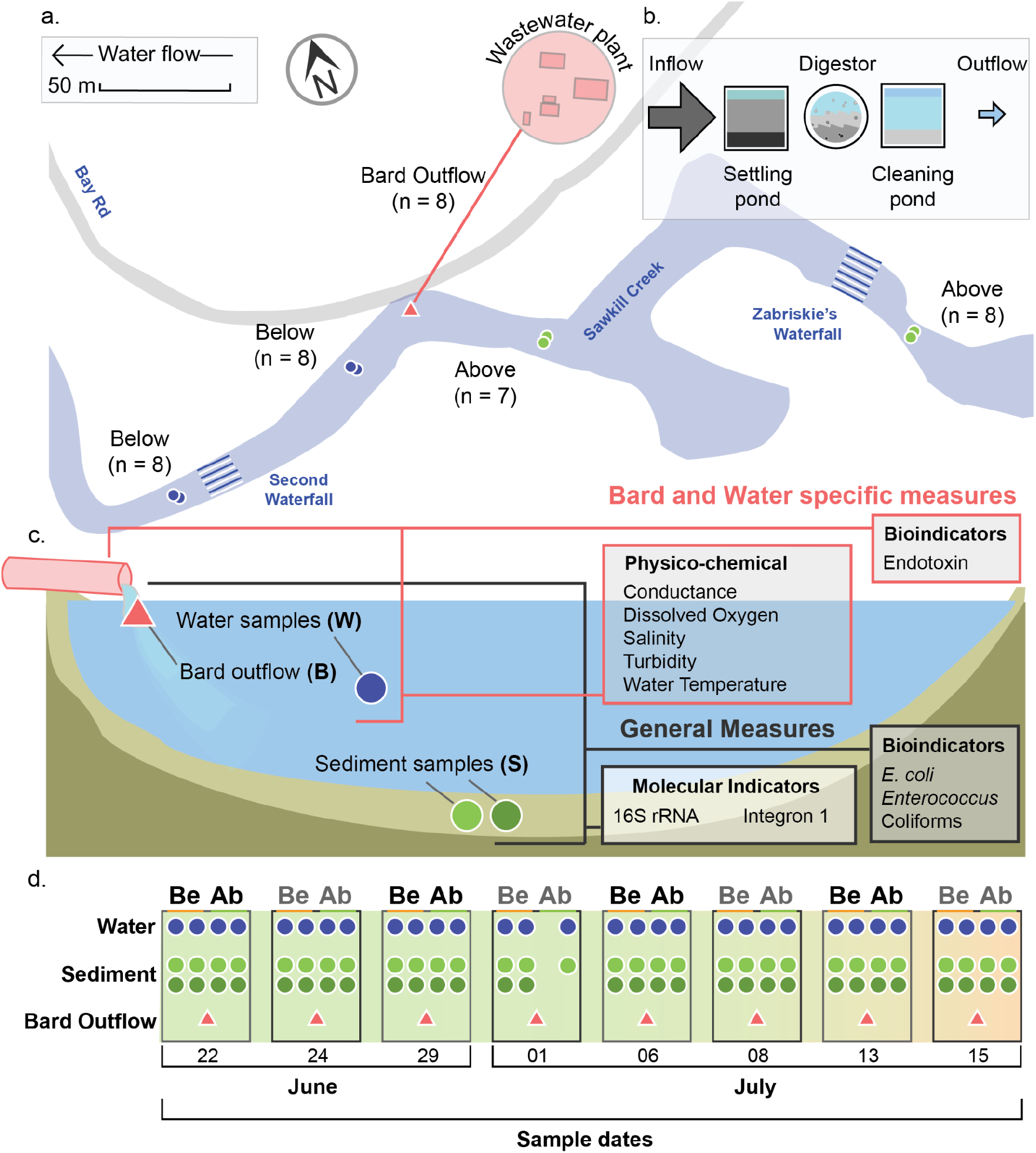
Diagram of study. Representative map of sampling sites used in this research with site names, total sample number (A). Schematic representation of the WWTP included in this study (B). Schematic representation of sampling types (Bard outflow (B); Water (W); Sediment (S)) and the sampling measures performed, including those applied to all samples (General Measures) and those specific to Bard and Water sample sites (C). Lastly, a timeline of 10 sampling events over 3 months shows the number of successful samples taken from the W, S, and B sample types for both the Below (Be) and Above (Ab) sites.

### 2.2 Water and sediment sampling

Our sampling, conducted during the summer of 2015, was focused on the Bard College (Annandale-on-Hudson, NY, USA) wastewater treatment plant (WWTP), which treats approximately 0.2 million gallons per day from the Bard College Campus (NYSPDES permit #’s NY0271420, NY0031925). In order to evaluate the specific contribution of WWTP effluent to the bacterial and genetic composition of water column and sediment communities near the outflow, we sampled at 2 sites above the outflow pipe (Fig. 1, Above) and 2 sites below the outflow pipe (Fig. 1, Below). We also sampled water from the outflow pipe itself to determine allochthonous inputs of bacteria and genetic material.

Surface water samples above and below the WWTP outflow were collected in sterile and acid-washed 2L Nalgene bottles from mid-channel within 0.5m of the stream surface at each site. Duplicate sediment cores (about 7 cm deep) were then taken from undisturbed sediments just upstream of the water sampling locations. The WWTP discharge was sampled using a sterile scoop and placed in a sterile and acid-washed 2L Nalgene bottle. All samples were placed on ice and transported to the laboratory for analysis within 2 hours of sampling. A list of all samples along with their physical characterization (i.e., GPS coordinates, date, temperature, salinity, conductance, particulate) is available in de Santana *et al*. (2022).

### 2.3 Sewage-indicator Concentrations

Using the IDEXX MPN method (IDEXX, Westbrook, ME, USA) with Enterolert and Colilert systems, we measured the concentrations of *Enterococcus, E. coli*, and Coliforms on both water and sediment samples within 2 hours of sample collection (ref). For Colilert assays, mid-channel water samples were diluted 1:10 with sterile DI water, and WWTP outflow samples were diluted at 1:10 and 1:100 with sterile DI water (refs). Sediment concentrations were acquired by creating a slurry sample by gently mixing 250 mg of centrifuged sediment with 50 mL of sterile DI water. A 1:10 and 1:100 dilution of the sediment slurry was then run through Enterolert and Colilert protocols (refs). According to manufacturer instructions, reagents for both Enterolert and Colilert IDEXX systems were gently mixed with sample water in a sterile 100 mL container. After full reagent dissolution, the contents were poured into a sterile 49-well Quanti-Tray (IDEXX, Wesbrook, ME, USA) and sealed. Colilert trays were then incubated for 24 hours at 35degC. Enterolert trays were incubated 24 hours at 41degC. Using chromatographic changes and fluorescent markers, concentrations or E. coli, Coliforms, and Enterococcus were calculated as MPN/ml. All data is further described here and available here ; SCI Data paper.

### 2.4 DNA Extraction

For each water sample, we filtered 750 mL through a 0.22 um Sterivex filter. We then extracted total DNA using the PowerWater DNA Isolation kit (MoBio Laboratories Carlsbad, CA, USA), now available as the DNeasy PowerWater DNA Isolation Kit (QIAGEN, Hilden, Germany). For each sediment sample, we also used the PowerSoil DNA Isolation Kit (MoBio Laboratories, Carlsbad, CA, USA), now available as the DNeasy PowerSoil DNA Isolation kit (QIAGEN, Hilden, Germany), to extract DNA from 250 mg of sediment.

### 2.5 *Quantifying* 16S rRNA *and* intI1 *genes*

To investigate the possible presence of molecular contaminants, we quantified the presence of the *16s rRNA* and *intI1* genes using quantitative PCR following the procedure described in de Santana et al (2022). While the *16S rRNA* gene encodes for the 16S subunit of the prokaryotic ribosome and is present in all known bacteria (10.1073/pnas.74.11.5088), *intI1* encodes the integrase component of the class 1 integron. The latter is a genetic mechanism mostly found in pathogenic bacteria such as *E. coli* and *Pseudomonas aerugnisa*, and can promote the horizontal gene transfer of antibiotic resistance gene cassettes (10.1186/s40168-018-0516-2). In brief, we processed each sample in triplicate using the primers described in Gaze et al (2011) and the PowerUp SYBR Green Master Mix (Applied Biosystems, Foster City, CA, USA). Samples were then ran using the Bio-Rad CFX96 Real-Time PCR Detection System (Bio-Rad Laboratories, Hercules, CA, USA) with an internal standard curve constructed from a serial dilution of the *E. coli* strain SK4903. Finally, we adjusted the total number of *16S rRNA* copies found in each sample by dividing by 4.2, which is the average number of *16S rRNA* copies each bacteria cell harbors^19^. Data files are available in de Santana et al. (2022).

### 2.4 16S rRNA *amplicon sequencing and analysis*

To characterize the overall bacteria community structure in each sample, we amplified the V4 region of our 16S gene amplicon sequencing library using primers 515F and 806R, as outlined in the Earth Microbiome Project^20^. Samples were sequenced on the Illumina Miseq platform using 250-bp paired ends by Wright Labs (Huntingdon, PA, USA) and are described in de Santana et al. (2022). Raw reads are publicly available on the Sequence Reads Archives of the NCBI (accession number: PRJNA565393). Resulting sequences were filtered and trimmed with Trimmomatic, ver. 0.39,21 using the following parameters: ILLUMINACLIP:TruSeq3-PE.fa:2:30:10 LEADING:3 TRAILING:3 SLIDINGWINDOW:4:15 M INLEN:100. Subsequent steps were performed on the Summer season subset of samples using QIIME2, ver 2021.2^22^ as described in de Santana *et al*. (2022).

DADA2 was used to denoise paired reads (--p-trunc-len-f 250, --p-trunc-len-r 233) with the median non-chimeric read count per sample 21,741 (Supplemental Table: Qiime2_Denoising_stats.tsv). Alpha-rarefaction (Faith-PD) suggests near complete representation for each site at approximately 13,000 reads (Supplemental Table: Supplemental_rarefaction_faith_pd.csv). Alpha-diversity between sites was determined using Shannon with down-sampling to 10,000 reads per sample (Supplemental Figure: Supplemental_Shannon_10K.pdf), we find that sites from the Sediment environmental source have the highest Shannon diversity, then the Outflow site, and finally the Water sites. There is no significant difference (Kruskal-Walis, p-value <= 0.05) between sites of the same environmental source (e.g. Water, Sediment, Outflow) although sites with different environmental sources are significantly different. We calculated beta-diversity using Weighted UniFrac down-sampled to 10,000 reads per sample (Supplemental Table: Supplemental_Beta-diversity_WU_10K.csv) and, similar to alpha-diversity, found the highest Distances occurred with the Sediment sites, then Outflow, and then Water. As with alpha-diversity, we found significant differences between the sites with different environmental sources (PERMANOVA, p-value <=0.05)

## 3. Results

### 3.1 Monitoring sewage-associated bacterial contaminants

#### 3.1.1 Sewage-associated bacterial contaminants in WWTP outflow

To determine whether the WWTP could represent a possible source of sewage contaminants in this system, we first investigated for the presence of viable *E. coli, Enterococcus* sp., and total coliforms in treated wastewater discharge. We detected the presence of sewage contaminants in only four samples out of the eight treated wastewater samples we investigated (Figure S1). Interestingly, when we did detect the presence of contaminants, their levels strongly correlated with the amount of rain events that happened in the previous 36 hours. In fact, the amount of precipitation directly resulted in a linear increase in *E. coli* concentrations (*F*_(1,6)_ = 28.64; *P* = 0.002; Figure 2A & S2A), *Enterococcus* sp. concentrations *F*_(1,6)_ = 18.63; *P* = 0.005; Figure 2B & S2B), and total coliforms concentration *F*_(1,6)_ = 8.01; *P* = 0.03; Figure 2C & S2C).

**Figure 2.**
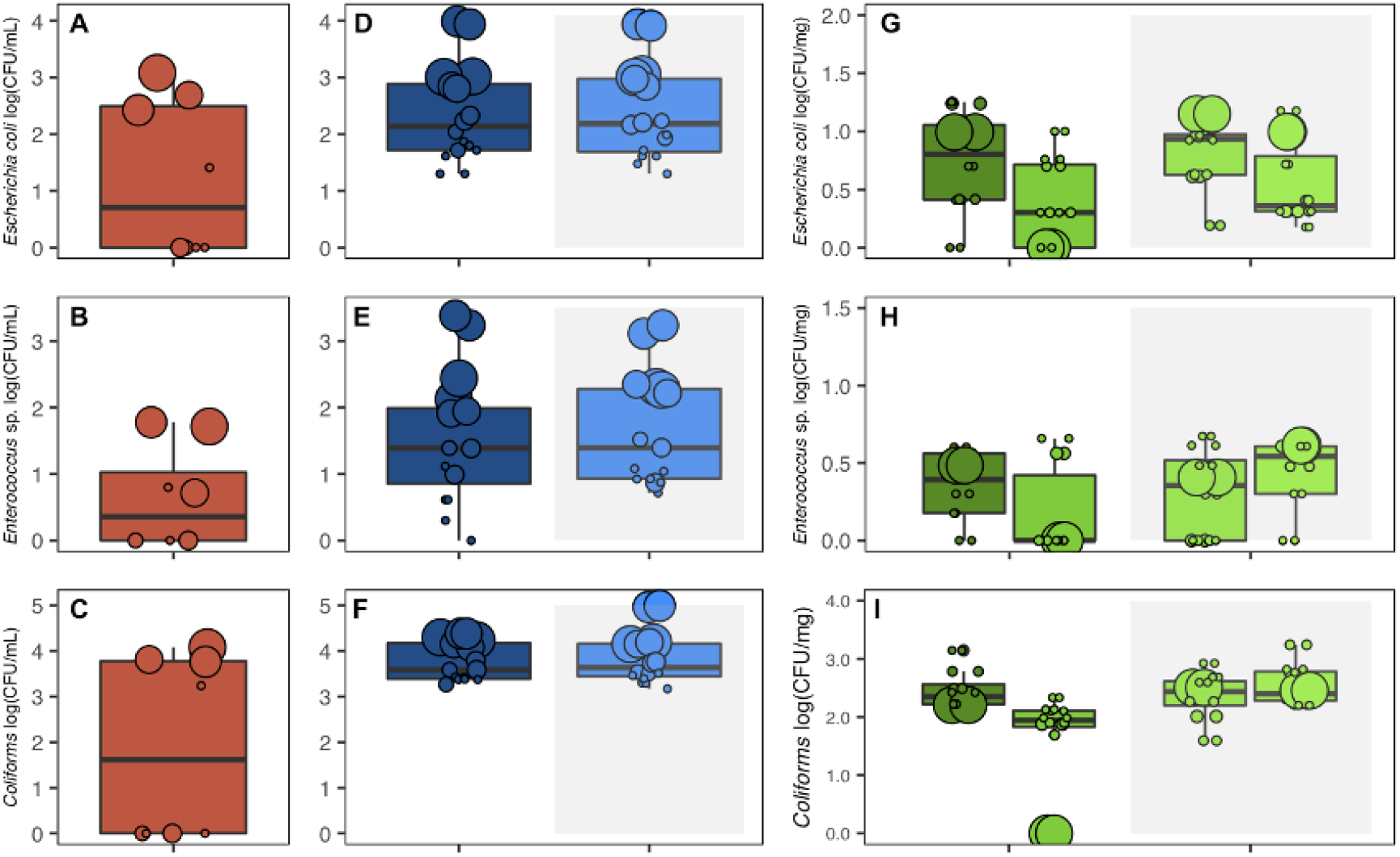
Shows the measured abundance of each sample of the outflow (**red**), the water samples from above the outflow (**dark blue**) and below (**light blue**), and the sediment samples from above (**dark green**) and below (**light green**). Abundances are for E. coli (**A, D, G**), Enterococcus (**B, E, H**), and Coliforms (**C, F, I**).

#### 3.1.2 Sewage-associated bacterial contaminants in surface waters

To determine whether the WWTP outflow was an important source of sewage contaminants in the environment, we then investigated the presence of *E. coli, Enterococcus* sp., and total Coliforms in two surface water samples above the outflow and two surface water samples below the outflow. After confirming that there was no statistical difference between the two samples taken at each site for all three indicators (Figure S2A-C), we found that there was no difference in the average concentration of *E. coli* (*F*_(1,29)_ = 0.03 ; *P* = 0.87; Figure 2 D), *Enterococcus* sp. (*F*_(1,29)_ = 039 ; *P* = 0.54; Figure 2 E), and total coliforms (*F*_(1,29)_ = 66.25 ; *P* < 0.0001; Figure 2F) above and below the outflow. Crucially, we found that the average concentration of *E. coli, Enterococcus* sp., and total coliforms were significantly lower in the outflow than both above (adj-*P*s < 0.0001) and below (adj-*P*s < 0.0001) water samples.

We also found that rain had a significant impact on the presence of all three sewage contaminants present in the surface water samples. More specifically, we found a positive linear relationship between the amount of rain in the previous 36 hours and the concentrations of *E. coli* (*F*_(1,29)_ = 62.49 ; *P* < 0.0001; Figure 2 D & S2D), *Enterococcus* sp. (*F*_(1,29)_ = 50.12 ; *P* <0.0001; Figure 2E & S2E), and total coliforms (*F*_(1,29)_ = 0.56 ; *P* = 0.46; Figure 2F & S2F). Interestingly, when we compared the average concentration of the sewage contaminant found in the outflow only during high rain events, we found that the concentrations were similar to that found in the above and below sites for all three sewage contaminants (adj-P > 0.05). In other words, during rain events, the concentration of *E. coli, Enterococcus* sp., and total coliforms were similar in the surface water and in the outflow. While our results suggest that the WWTP can be a source of sewage contaminants during high rain events, our results suggest that the main source of contamination is external to the WWTP.

#### 3.1.3 Sewage-associated bacterial contaminants in sediments

We then investigated whether sediment could act as a reservoir of sewage contaminants by characterizing the presence of *E. coli, Enterococcus* sp., and total coliforms in sediments collected at two sites located above the outflow and two sites located below the outflow. We detected the presence of all three sewage contaminants (Figure 2G-I), but unlike the surface water, we found that the samples’ specific site within each location significantly affected the concentration of *E. coli* (*F*_(2,59)_ = 6.52 ; *P* = 0.003; Figure 2G) and total coliforms (*F*_(2,59)_ = 10.61 ; *P* = 0.0001; Figure 2I). More specifically, we found that the concentration of both *E. coli* and total coliforms was the lowest in sediment samples from the site located just above the outflow (adj-*P*s < 0.001). While this result could suggest that the concentrations of *E. coli* and total coliform are lower above the WWTP outflow, it is important to note that we did not find a statistical difference between the levels of *E. coli* or total coliforms between the sites located below the outflow and the upper site located above the outflow (adj-Ps > 0.41).

Also, we found that rain events affected the distribution of sewage contaminants in the sediments (Figure 2G-I; Figure S3). Overall, we found that increased rain events in the previous 36 hours positively correlated with an increase in the concentration of *E. coli* (*F*_(1,59)_ = 10.81 ; *P* = 0.002; Figure 2G & S2G) and *Enterococcus* (*F*_(1,59)_ = 13.98 ; *P* = 0.0004; Figure 2H & S2H) found in the sediments. Interestingly, while we observed that the largest rain events led to the highest *E. coli* and *Enterococcus* sp. concentrations at most sampled sites, we also observed that the same rain events resulted in concentrations below the detection levels for the two contaminants at the site just above the outflow. The latter indicates that the presence of live sewage contaminants in sediments likely depend on the local geo-physical conditions of any sites, with varying sedimentation and resuspension rates. Finally, we found that an increase in rain precipitation had an overall negative effect on the concentration of total coliforms (*F*_(1,58)_ = 6.37 ; *P* = 0.01; Figure S2I), suggesting that the Coliforms might be more easily resuspended from sediments during rain events.

### 3.3 Monitoring molecular contaminants: 16S rRNA gene and integron 1 gene

#### 3.3.1 16S rRNA *genes in WWTP, surface waters, and sediments*

While WWTP processes are designed to minimize the release of live sewage contaminants, in most instances WWTP are not required to monitor the release of molecular contaminants such as DNA. To investigate whether WWTP outflow could be a source of molecular contamination, we monitored the abundance of the *16S rRNA* gene and the *intI1* gene in the outflow as well as in surface water and sediment samples. While *16S rRNA* is an indicator of the overall quantity of bacterial DNA being released in the environment, the *intI1* gene is an indicator of the likely presence of antimicrobial resistance in Gram-negative bacteria.

As expected, we found a high abundance of *16S rRNA* genes in all samples, including the WWTP outflow (Figure 3A), surface waters (Figure 3B), and the sediments (Figure 3C). Interestingly, while we found that the abundance of 16S rRNA copies was similar in the outflow and in all of the surface water samples (adj-*P*s > 0.05), we found that 16S rRNA abundance was on average higher below the outflow than it was above the outflow (*F*_(1, 27)_ = 11.22 ; *P* = 0.002; Figure 3B). Moreover, we found that the abundance of *16S rRNA* copies was significantly shaped by the precipitation events in the 24 hours before the measures (Rain24:Position: *F*_(1, 27)_ = 8.16; *P* = 0.008; Figure 3B). More specifically, we found that, overall, *16S rRNA* copies tend to decrease with higher rain precipitation, but that this trend was stronger above the outflow (Figure S5B). Finally, we found that *16S rRNA* abundance in sediments did not differ between sites, above or below the outflow, but were again negatively impacted by the amount of precipitation events up to 36 hours preceding sampling (*F*_(1, 54)_ = 7.13; *P* = 0.01; Figure S5C).

**Figure 3.**
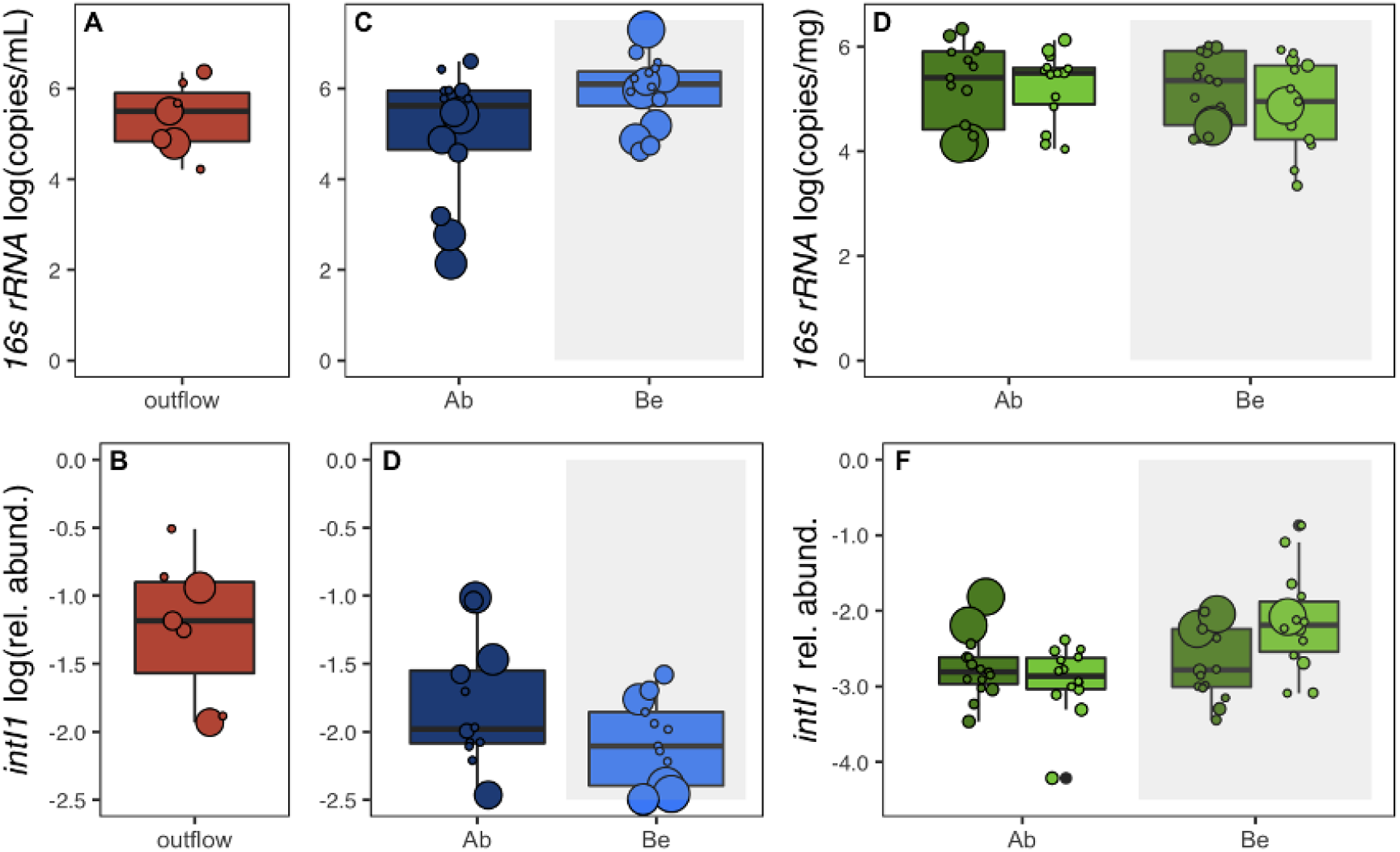
Shows the measured abundance of each sample of the outflow (**red**), the water samples from above the outflow (**dark blue**) and below (**light blue**), and the sediment samples from above (**dark green**) and below (**light green**). Abundances are for *16s rRNA* (**A, C, E**) and IntI1 (**B, D, F**).

#### 3.3.1 Integron 1 gene in WWTP, surface waters, and sediments

We further characterized the possible impact of the molecular contaminants on the surrounding environments by estimating the abundance of the *intI1* gene, an important indicator of human disturbance, relative to the abundance of *16S rRNA* gene.

Interestingly, we found that *intI1*’s relative abundance in the outflow (Figure 3D) was significantly higher than it was in the samples above (t= ; *P* < 0.0001) and the samples below the outflow (t = ; *P* < 0.0001), but did not differ between sites below and above the outflow (*F*_(1, 28)_ = 1.88 ; *P* = 0.18 ; Figure 3E). We found no statistically significant differences in *IntI1* concentrations relative to rain events.

Finally, while we found the presence of *intI1* in all sediment samples, we found that its relative abundance was higher in sediments located below the outflow (*F*_(2, 54)_ = 4.51 ; *P* = 0.01 ; Figure 3F). More specifically, we found that *intI1*’s relative abundance was higher in the site directly below the outflow than it was at any other site (min adj-*P* < 0.03).

### 3.4 The effect of WWTP on bacterial communities

#### 3.4.1 Bacterial communities in the WWTP

To investigate whether the WWTP outflow had a significant effect on the bacterial communities present in the local environments, we studied the overall bacterial diversity present in the outflow and at sites below and above the outflow using *16S rRNA* amplicon sequencing. From 97 samples, we sequenced a total of 2,257,390 (post-quality control) paired reads (Supplemental table-dada2.qsv) that were assigned to 12,138 amplicon sequence variants, or ASVs (Table S1_ASV_to_Taxa.tsv). We identified 3001 distinct taxa composed of 70 different phyla, 415 orders, and 1007 genera among the ASVs (Table S1_ASV_to_Taxa.tsv). While a full list of taxa is available (Table S1_ASV_to_Taxa.tsv), the most common phyla were *Proteobacteria* (50.4% ± 9.98) followed by *Bacteroidota* (15.9 ± 11.9%), *Actinobacteriota* (7.8 ± 9.6%), and *Acidobacteriota* (4.7 ± 4.2%). *Comamonadaceae* (14.6 ±10.9%), was the most frequent family followed by *TRA3-20* (order *Burkholderiales)* (4.8 ± 4.3%), *Sporichthyaceae* (4.7 ± 7.1%), *Spirosomaceae* (3.9 ± 5.8%), *Chitinophagaceae* (3.8 ± 3.2%), and *Rhizobiales_Incertae_Sedis* (3.6 ± 3.2%)(Figure S2A). Only a small minority of ASVs (319, 2.6%) were found in all three ecological niches, we found that 2018 (16.6%) ASV were present in surfaces waters exclusively, 7293 (60.1%) ASVs in sediments only, and 1207 (9.9%) were ASVs in the outflow only.

Because of the relevance of the WWTP outflow microbiome to the experiment, we consider it here in greater detail. We found a total of 2566 ASVs identified as part of 40 different phyla, 250 orders, and 378 genera. More specifically, the most common phyla were Proteobacteria (61.8% ± 14.7) followed by Bacteroidota(15.9 ± 7.3%), Actinobacteriota(6.0 ± 5.8%), and Campilobacterota (3.4 ± 4.5%).

*Comamonadaceae*(25.6 ± 12.4%), was the most frequent family followed by *Rhodocyclaceae* (6.4 ± 4.9%), *Neisseriaceae* (4.8 ± 5.7%), *Flavobacteriaceae* (4.8 ± 3.2%), *Arcobacteraceae* (3.3 ± 4.4%), and *Aquaspirillaceae* (2.5 ± 3.9%)(Figure S2A). Interestingly, while we found substantial concentrations of bacterial contaminants such as *E. coli* and *Enterococcus* spp. in the WWTP outflow, we did not find their molecular signature in the outflow. This suggests that the bacterial contaminants are found in relatively low abundance compared to other bacteria types and indicate that it is important to consider the broader diversity found in microbial communities to understand the possible impact of WWTP on local communities.

#### 3.4.2 Bacterial communities in surface waters around the WWTP

Because the WWTP outflow discharges directly into the water, we first investigated the possible impact of the outflow on microbial communities found in surface waters. More specifically, while the number of ASV observed in the outflow was significantly lower than in surface water above or below the outflow (KW test on Observed: *P* = 0.001496), we found that the number of observed ASVs did not different between microbial communities samples above and below the outflow (KW test on Observed: *P* = 0.782). Similarly, we found that prokaryotic diversity, measured as the Shannon index, was also higher in the WWTP (Supplemental_Shannon_10K.pdf) outflow when compared to communities samples from above and below surface waters (KW test on Shannon: p-value = 4.0e-3). Again, we found that the ASV diversity was not different in prokaryotic communities found in the water samples above or below the outflow (KW test on Shannon: p-value = 0.8587).

We then investigated whether the bacterial community structure changed between the different sites using pairwise comparison of genera abundance (DESeq2, Wald, adj-*P* <= 0.05). From all the comparisons, we found that the greatest difference in microbial communities was observed between the Above and Outflow sites, with 16 genera being significantly higher in the Outflow and two genera higher in the Above sites (Table SB: Deseq2_sig_diff_ Outflow_v_ Above_(far_near)_water_Summer_ wald.csv). Interestingly, we found a single genus (*Macromonas*) changed in abundance between communities found Above and Below the outflow (Table S: Deseq2_sig_diff_Below_v_Above_(far_near)_water_Summer_wald.csv) and that no genera significantly changed in abundance between the communities found in the Outflow and Below the outflow.

#### 3.4.3 Bacterial communities in sediments around the WWTP

We then investigated the possible impact of the outflow on microbial communities found in sediments by comparing bacterial diversity found Above and Below the outflow. Overall, we found that prokaryotic diversity was higher in all sediment samples relative to the outflow (KW test on Shannon diversity, p-value = 8.1e-4). More specifically, while the number of ASV observed in the outflow was significantly lower than in surface water above or below the outflow (KW test on Observed: *P* = 0.001496), we found that the number of observed ASVs did not differ between microbial communities samples above and below the outflow (KW test on Observed: *P* = 0.782). Similarly, we found that prokaryotic diversity, as measured by the Shannon index, was also lower in the WWTP outflow when compared to sediment communities samples from above and below surface waters (KW test on Shannon: p-value = 0.0073). Again, we found that the ASV diversity was not different in bacterial communities found above or below the outflow (KW test on Shannon: p-value = 0.8587).

We then investigated whether the bacterial community structure changed between the different sites using pairwise comparison of genera abundance (DESeq2, Wald, adj-*P* <= 0.05). While we find a single genus, *Syntrophorhabdus*, significantly different between Below and Above bacterial communities, we found that the bacteria communities found in sediments differed greatly compared to the outflow communities. More specifically, we found that 65 genera are significantly different between Outflow and Below site sediment, with 30 being higher in the Outflow sediment and 35 higher in the Below sediment (Table SA: Deseq2_sig_diff_Outflow_v_Below_(far_near)_sediment_Summer_wald.csv). Similarly, we found 88 genera are significantly different between the Above and the Outflow sites, with 56 being higher in communities found Above and 32 higher in the communities found in Outflow (Table SC: Deseq2_sig_diff_Outflow_v_Above_(far_near)_sediment_Summer_wald.csv). These results suggest that the WWTP outflow has a limited impact on bacteria communities found in sediment when measured at the genus level.

### 3.5. The impact of temporal change and precipitation on bacterial communities

#### 3.5.1 Temporal changes in bacterial communities

To investigate the possible role of environmental variables in shaping the bacterial communities in relation to sampling sites, we used linear models to identify taxa with differential correlation in regard to specific variables of interest. More specifically, using Limma differential abundance analyses of longitudinal data, we identified all the taxa with a statistically significant difference relative to the background and with a minimal abundance in the population.

Taking the genus as the lowest taxonomic level of resolution, we found 14 taxa (out of 781) from the Water samples that significantly varied from the global background over time ((*P* < 0.05, minimum abundance 0.1%; Figure 4A). Similarly, a longitudinal analysis of the Sediment samples identified 4 taxa (out of 1023) that were changed significantly, relative to the background, over time (*P* < 0.05, minimum abundance 0.05%; Figure 4B). From both sample sets there is a pronounced shift in population structure occurring on the day of July 1, 2015, corresponding to the large rain event described later.

**Figure 4.**
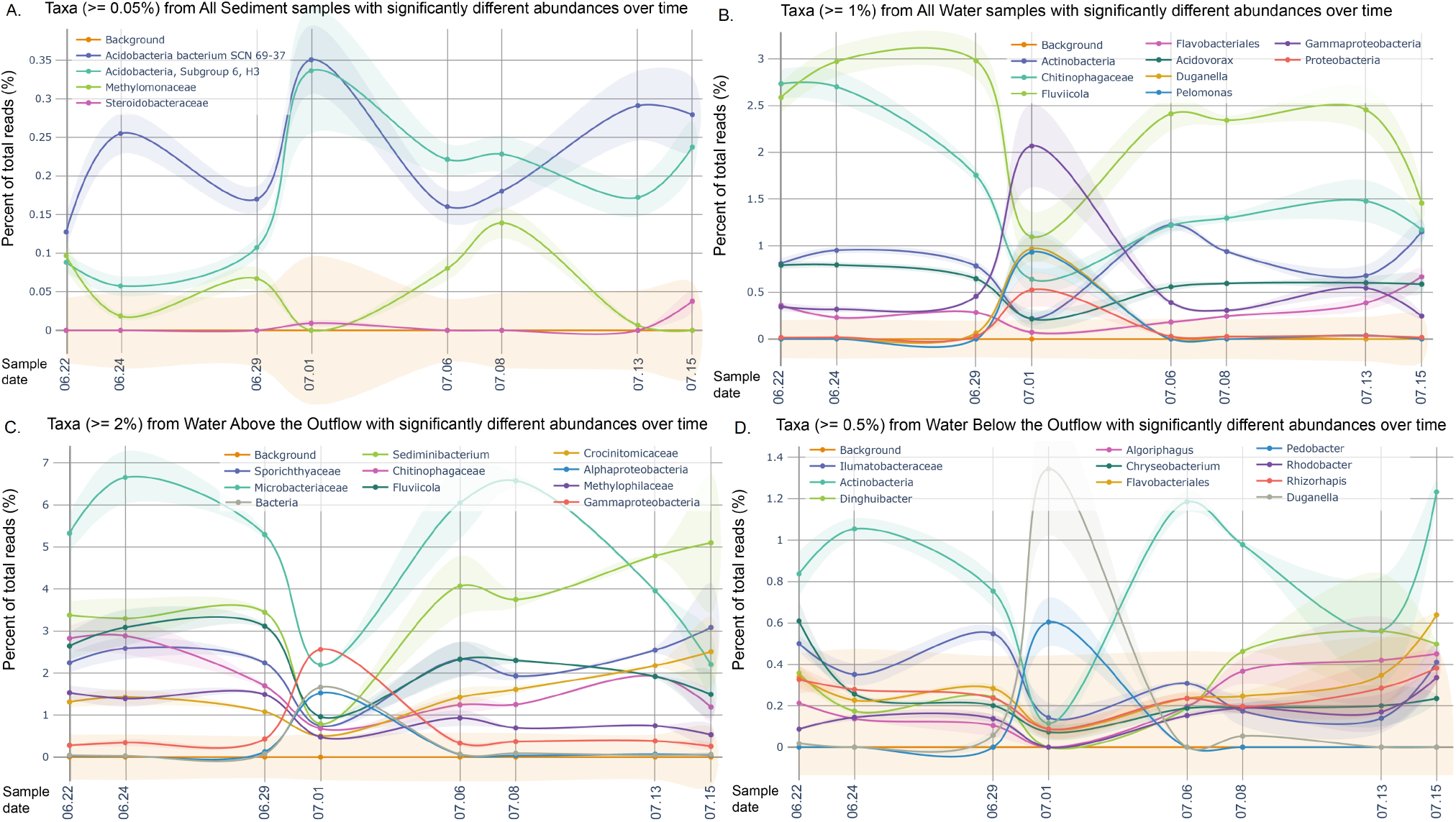
Taxa with significantly different relative abundances over time relative to background. Here, we show the results of a longitudinal analysis of changes in relative taxa abundance over time. Plotted lines are those taxa significantly different from the background (Limma, p-value <= 0.05) with the 1 standard deviation for the taxa shown as the translucent band. The Sediment samples (**A**) had fewer taxa (5) with minimum relative abundance (>= 0.05%) than that observed for the Water samples (B, C, D). The Water samples, combining both Above and Below sites (**B**) had nine significantly different taxa, with a high minimum relative abundance (>= 1%). The Water samples are further separated into the Above class (**B**, 10 taxa, >= 2%) and Below class (**D**, 10 taxa, >= 0.5%).

Separating samples based on whether they were above or below the Bard Outflow reveals different taxa varying over time and in response to the rain event. Water samples from above the Outflow (*P* < 0.05, minimum abundance 2%; Figure 4C) had larger changes in higher abundance taxa relative to the below Outflow samples (*P* < 0.05, minimum abundance 0.5%; Figure 4D). Crucially, we found some taxa that changed in abundance in the water bacterial communities and also changed in abundance in the Outflow communities. More specifically, we found Gammaproteobacteria, a bacterial class that include many bacterial contaminants of interest such as *E. coli*, as well as *Duganella* sp. and *Pelomonas* sp., two bacteria genus associated with human activity, among other all increased in abundance by at least a two-fold or more. On the other hand, *Fluviicola* sp., a genus of freshwater bacteria and one of the most abundant taxa in our water samples, decreased by more than a two-fold in abundance at the same time. Furthermore, when we analyze bacterial communities from Above and Below sites independently, we found that different ASVs change in abundance on the July 1, 2015 event. Namely, we found that Gammaproteobacteria undergo the largest increase in abundance in Above communities (Figure SA) and that *Duganella* sp. and *Pedobacter* sp. see that large abundance increases in Below communities (Figure SB). These results suggest that an increase in Gammaproteobacteria is more likely to result from upstream activity and that the Outflow likely produces the two bacteria names above.

#### 3.5.2 Precipitation and changes in bacterial communities in water

Based on available environmental data (see methods), we predicted that precipitation, or rain, is the main environmental factor responsible for bacterial community shifts observed on July 1, 2015. Indeed, we found that 14 bacterial genera significantly changes in abundance in the Outflow in correlation with the amount of rain observed in 24 hours observed prior to sample collection (P < 0.05; Figure 5A). Similarly, we observed that rain had a significant impact on surface waters, with eight taxa varying distinctly in proportion to the rain (Figure 5B). Interestingly, when we consider bacterial communities from Above and Below waters independently, we found that different ASVs are enriched. While we see an increase in abundance in *Pseudomonas, Enterobacteriaceae, Rickettsiella* among others, above the Outflow (Figure SA), we only see a substantial increase in *Pseudomonas* sp. below the outflow (Figure SB). These results suggest that the presence of *Pseudomonas* and *Enterobacteria* in the near environment of the WWTP is likely to be explained by an upstream source.

**Figure 5.**
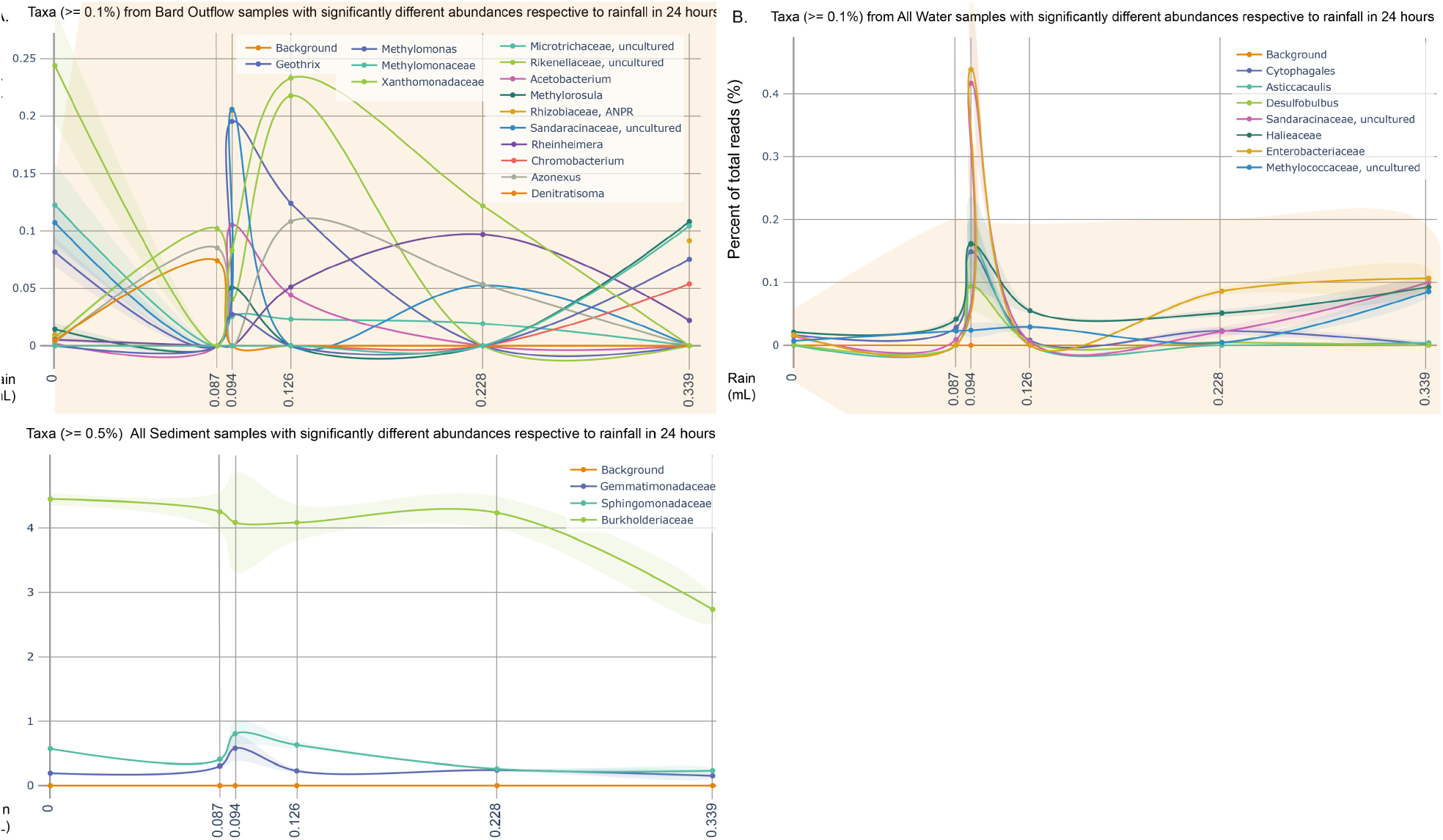
Taxa with significantly different relative abundances in relation to rainfall (24 hrs) relative to background. Here, we show the results of an analysis of changes in relative taxa abundance in relation to rainfall in the preceding 24 hours. Plotted lines are those taxa significantly different from the background (Limma, p-value <= 0.05) with the 1 standard deviation for the taxa shown as the translucent band. The Outflow samples (**A**) had the highest number of taxa (14) with minimum relative abundance (>= 0.1%) than that observed for the Water (**B**) or Sediment (**C**) samples. The Water samples (**B**), combining both Above and Below sites (**B**) had 7 significantly different taxa. While the Sediment samples (**C**) had the least number of significantly different taxa (3).

We further investigated further the possible role of a single heavy rain event on bacterial communities by comparing ASVs abundance before the heavy rain event (i.e., June 22, 2015, June 24, 2015, and June 29, 2015) to that of July 1, 2015 immediately after the heavy rain event of June 30th, 20215. Using DESeq2, we found that three genera significantly decrease in abundance in the Outflow communities following the heavy rain event (Table SA), and that 31 genera also significantly decreased in the surface waters above the Outflow (Table SB), and 12 genera decreased in surface waters below the Outflow (Table SC). These results suggest that while precipitation can increase the relative abundance of certain taxa, the main effect of large precipitation events is to dilute the abundance of most bacteria genera in the outflow and in surface waters.

#### 3.5.3 Precipitation and changes in bacterial communities in sediment

Finally, we investigated how precipitation events impact bacterial communities found in sediments above and below the Outflow. Using linear models to identify ASVs that may correlate with the intensity of precipitation events, we found only three bacterial classes to change in relative abundance in response to the main precipitation event (Figure 5C). However, when we considered bacteria communities from Above and Below independently from each other, we observed major differences. This time, we found that 10 taxa were significantly different from the Above samples (>=0.1%), while 50 were observed for the Below samples (Figure SA and Figure SB), with only two taxa shared between both sets. While the Below samples evidenced higher numbers of significantly different taxa these could not be shown to be exclusively a product of contamination from Bard Outflow, as each taxa was also identified in the Above sediment samples, although often at much reduced relative abundance.

Again, we further investigated the possible role of a single heavy rain event on bacterial communities by comparing ASVs abundance before the heavy rain event (i.e., June 22, 2015, June 24, 2015, and June 29, 2015) to that of July 1, 2015 immediately after the heavy rain event of June 30th, 20215. Using DESeq2, we found no significant change in terms of abundance between the Above and Below bacterial communities. These results suggest that rain events likely happen often enough that all soil looks disturbed and could potentially act as a source of bacterial contamination in the surface waters.

## 4. Discussion

In this study we assessed the possible effects of a wastewater treatment plant (WWTP) in a freshwater environment, by comparing samples from the outflow with sediments and water samples collected from above and below the WWTP. The samples were analyzed for fecal indicator bacteria, molecular contamination and prokaryotic communities along the temporal and longitudinal lines, while addressing the effects of rain events in the recorded weather patterns. Overall, we found that the fecal-indicator bacteria were mostly correlated with rain events and that the WWTP is efficient in eliminating these organisms. However, while we did not find evidence of living fecal indicator bacteria in the WWTP outflow, we did find substantial abundances of *16S rRNA* genes associated bacteria, as well as *IntI1*, a known driver of horizontal gene transfer (HGT) of antibiotic resistant genes (ARGs), suggesting the WWTP is not as efficient in eliminating molecular contamination.

More specifically, the higher concentration of sewage-associated bacteria in the waters from above and below the outflow in comparison to the discharge of the WWTP is suggestive that there are sources of sewage contaminants located upstream of the WWTP outflow. The rain event in the previous 36 hrs caused significant changes in sewage-associated bacteria in the sediments as well as observed for water samples. The effects of rain events on sewage-associated bacteria can last for several days depending on the local conditions (Mueller-Spitz et al., 2010).

Interestingly, we found that the concentration of *intI1* in the outflow or in the surface water samples was not as affected by rain event as it was the case for the raw sewage contaminant, suggesting that the outflow release of molecular contaminant is a process not related to the release of raw sewage observed during high rain event. For the sediments, however, higher concentrations of *IntI1* were found in the below site closest to the WWTP, suggesting that the outflow is a potential source of molecular contamination linked to antimicrobial resistance, as previously suggested (Young et al., 2013) and that sediments can act as a reservoir where certain genes can accumulate under environmental pressures, which has been confirmed (Mao et al., 2013 [10.1021/es404280v]).

We also analyzed the different sample types in regards to the overall prokaryotic communities. The sample types showed little overlap of ASV indicating that there are important differences in the communities at each matrix. Interestingly, the sewage-associated taxa observed in the specific analysis were not found in the analysis of the entire prokaryotic communities. This suggests that the bacterial contaminants are found in relatively low abundance compared to other bacteria types and indicates that it is important to consider the broader diversity found in microbial communities to understand the possible impact of WWTP on local communities.

When considering the *16S rRNA* gene as a proxy for prokaryotic diversity, we found the sediments to be more diverse as expected according to the literature (Jiang et al., 2006; Zeglin, 2015), while the outflow samples showed a higher diversity (Shannon) in comparison to the water samples. It is important to point out that as previously mentioned, the WWTP does not seem to efficiently eliminate molecular contaminants, and thus, there is a possibility that part of the microbial diversity addressed by the *16S rRNA* gene is due to free DNA released from the outflow rather than actual organisms. That said, the viability of free DNA in the environment is relatively unknown given the variety of degradation rates, which are influenced by a variety of environmental factors (Yang et al., 2020 [10.1016/j.scitotenv.2020.140592]). Therefore, while some part of the diversity found can be artificial, we consider that it would not be relevant. Considering the taxonomy, the water samples collected below the outflow showed more similar taxa with both waters from above and the outflow, while the outflow showed lower similarity with taxa from upstream. These results suggest that the bacterial communities found below the outflow are both influenced by bacterial communities coming from upstream as well as the WWTP outflow. The influence of the outflow in the sediments appears to be lower, with the sediments from above and below the WWTP displaying similar diversity and taxonomic profiles.

Notably, our sample collection included a moderately sized, once in a decade by volume, rainfall event on July 1, 2015. This event had a pronounced impact on species diversity and composition across all sites including sediment samples. This perturbation could have been caused by runoff from the creek banks, managed overflow from the WWTP, disturbance of the sediment (churn), or a combination of these factors. While previous literature has evaluated the role of precipitation in microbial community composition, this work suggests that even moderate rainfall events themselves may play a role in large scale re-ordering of microbial population structures. Indeed, the interactions between wastewater and stream communities with environmental factors is very complex (Burdon et al., 2020). This is particularly notable given the expected increase in large scale local weather events due to global climate change.

## Abbreviations

FIB: fecal indicator bacteria
WWTP: wastewater treatment plant
CSO: combined sewer overflow
SPDES: State Pollutant Discharge Elimination System

## Acknowledgements

This work was funded by Bard College and the Bard Summer Research Institute of Bard College. We thank Alexandra Clarke, Haley Goss-Holmes, Pola Kuhn, Tierney Weymueller, Marco Spodek, Christopher Hulbert, Beckett Landsbury, and Yuejiao Wan for technical assistance in the lab.

## Contributions

Work was planned by ED and GGP DA was associated with collection of the samples and contributed to the DNA sequencing. COS and PS worked on raw data analysis and the draft of the article with ED and GGP. GGP made final revisions on the manuscript.

## Corresponding author

Correspondence to Gabriel G. Perron

## Supplemental Materials

**Figure S1.**
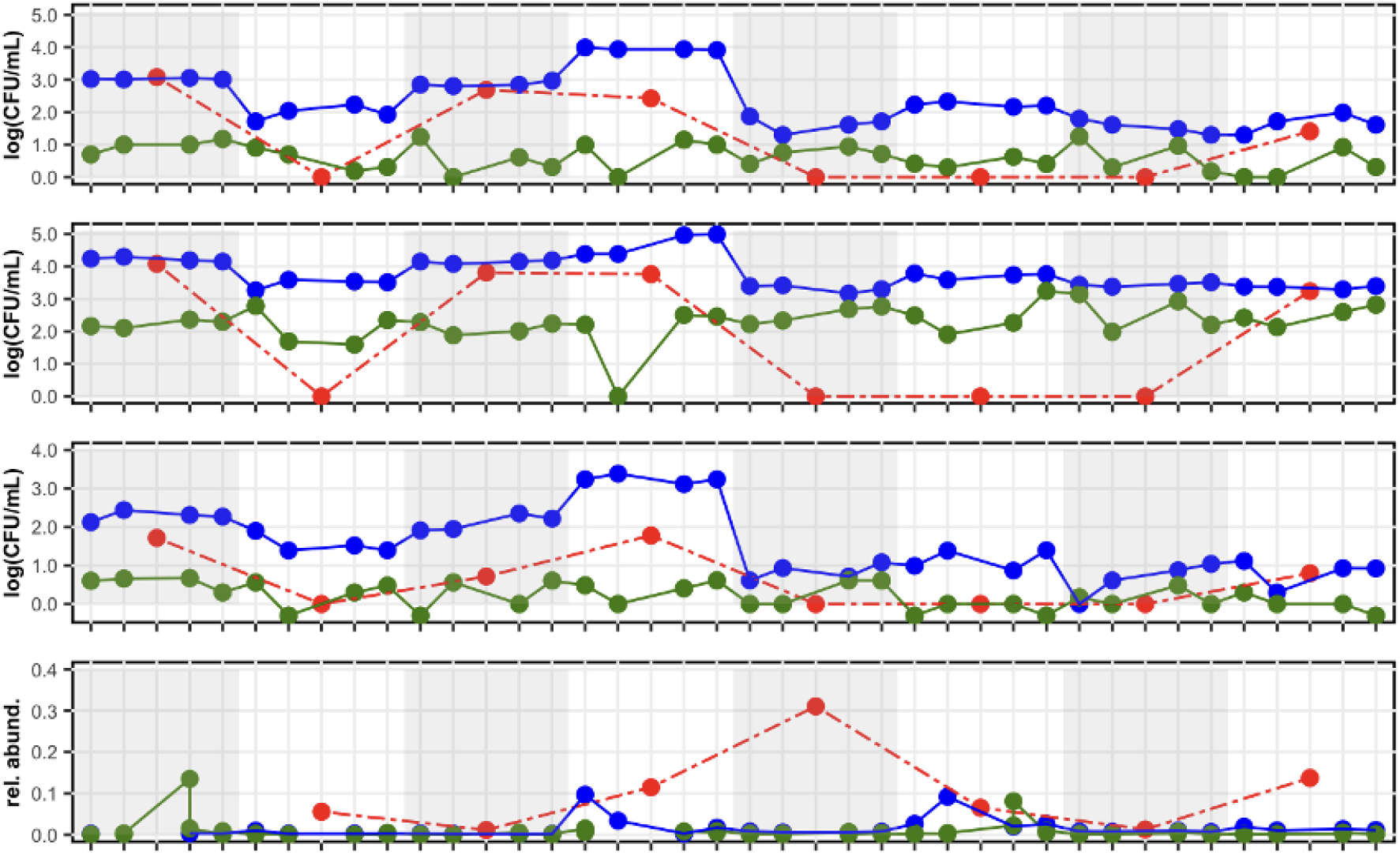
Longitudinal measurements of three sewage contaminants and one molecular contaminant. For each collection date, samples were taken from the WWTP outflow discharge and at two sites above and two sites below the outflow and were evaluated for: a) *E. coli* concentration (CFU/mL in water and CFU/mg in sediment); b) Coliform concentration (CFU/mL and CFU/mg in sediment); c) *Enterococcus* sp. concentration (CFU/mL and CFU/mg in sediment); and d) integron 1 relative abundance measured as the copy number of *intI1* gene over the copy number of the *16S rRNA* gene. Outflow and water data points depict the raw value for each sample while sediments data points (green) are the average of two biological replicates. Surface water samples are presented in blue, sediments in green, and WWTP outflow in red.

**Figure S3.**
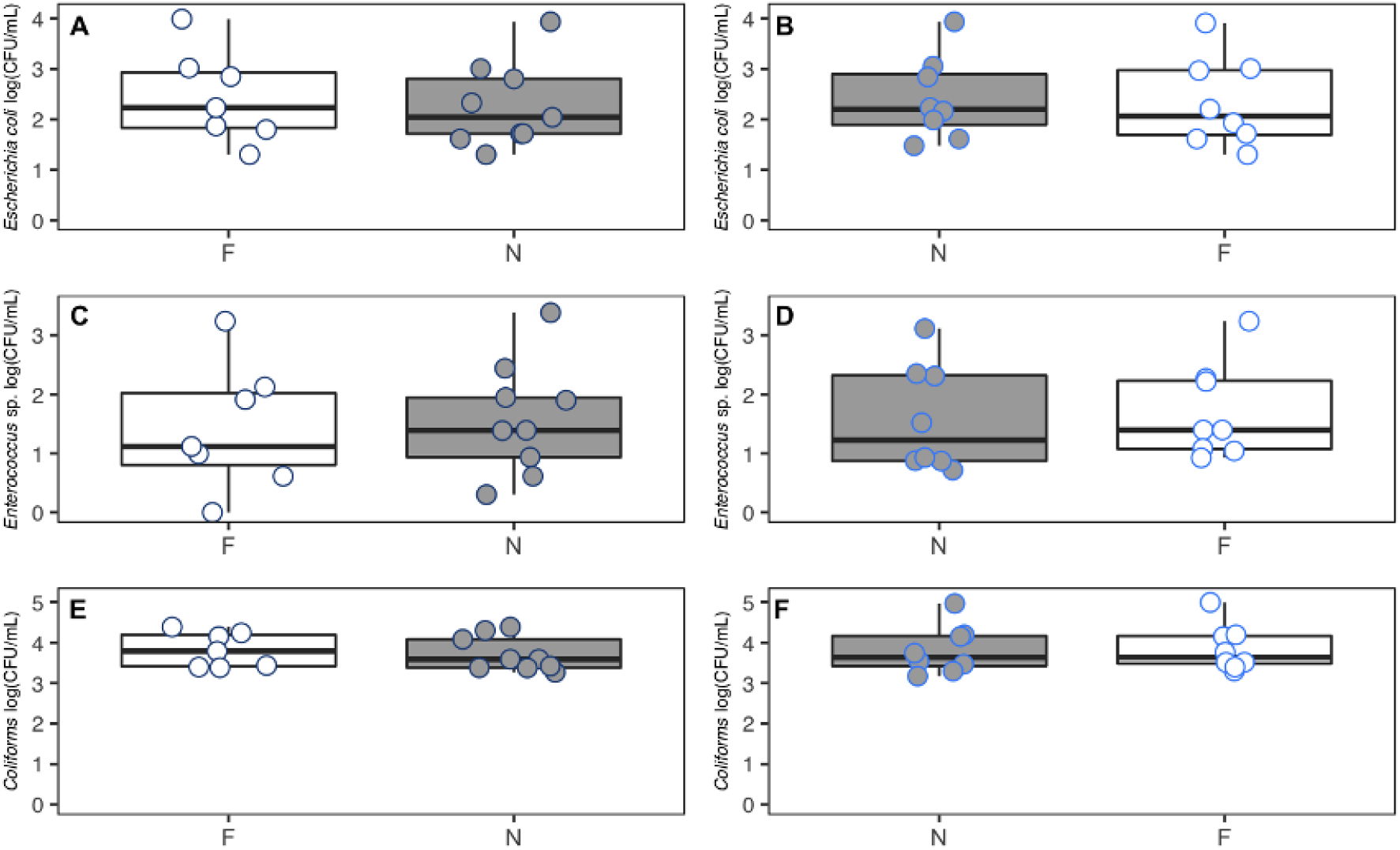
Concentration of sewage contaminants in two sites within locations. In outflow for A) *E. coli*; B) *Enterococcus* sp.; and C) total coliforms. In surface water for A) *E. coli*; B) *Enterococcus* sp.; and C) total coliforms. In sediments for A) *E. coli*; B) *Enterococcus* sp.; and C) total coliforms.

**Figure S4.**
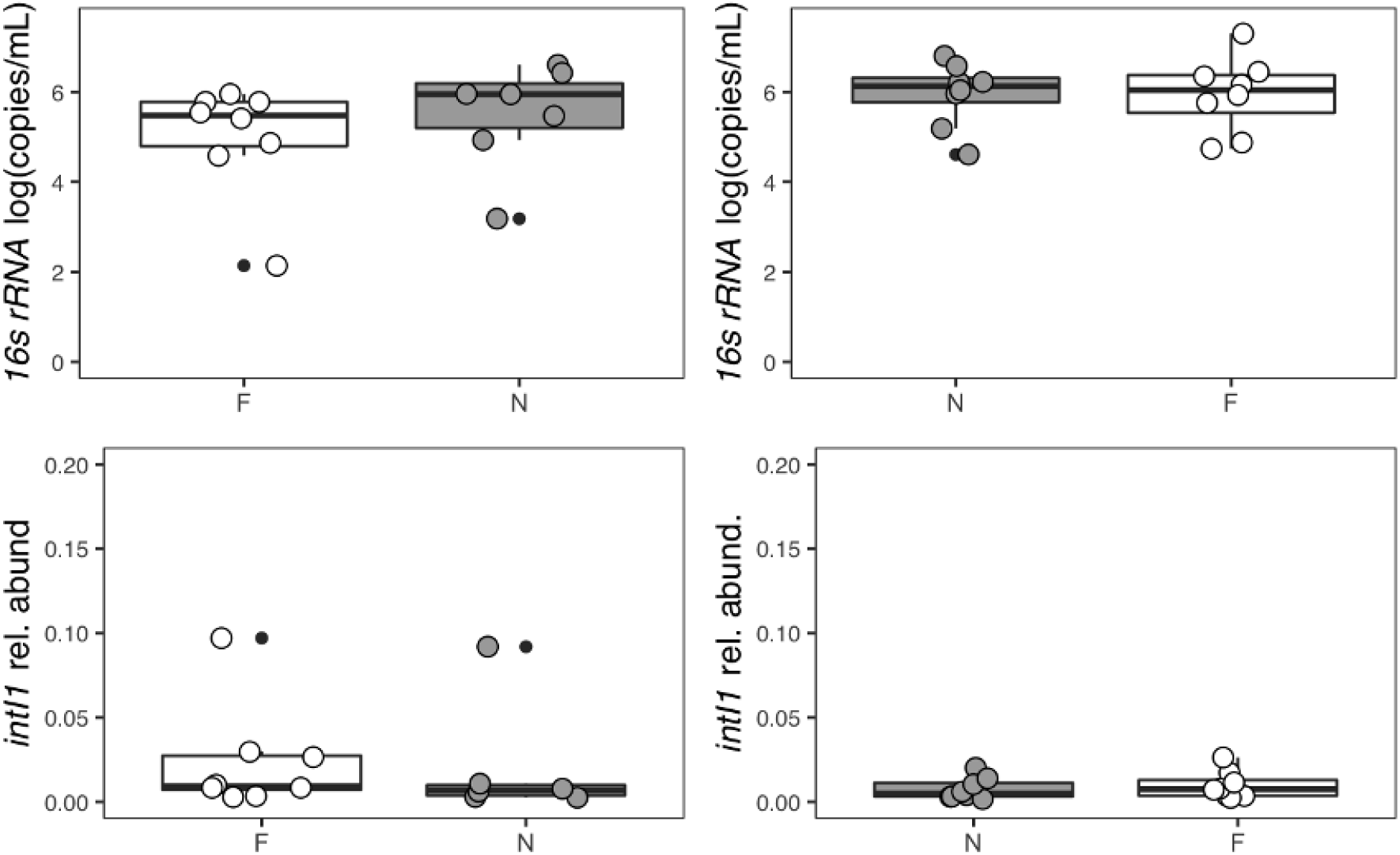
Concentration of molecular contaminants at two sites within locations. In outflow for A) *16SS rRNA* B) *intI1* sp.; and C) total coliforms. In surface water for A) *E. coli*; B) *Enterococcus* sp.; and C) total coliforms. In sediments for A) *E. coli*; B) *Enterococcus* sp.; and C) total coliforms.

**Figure S5.**
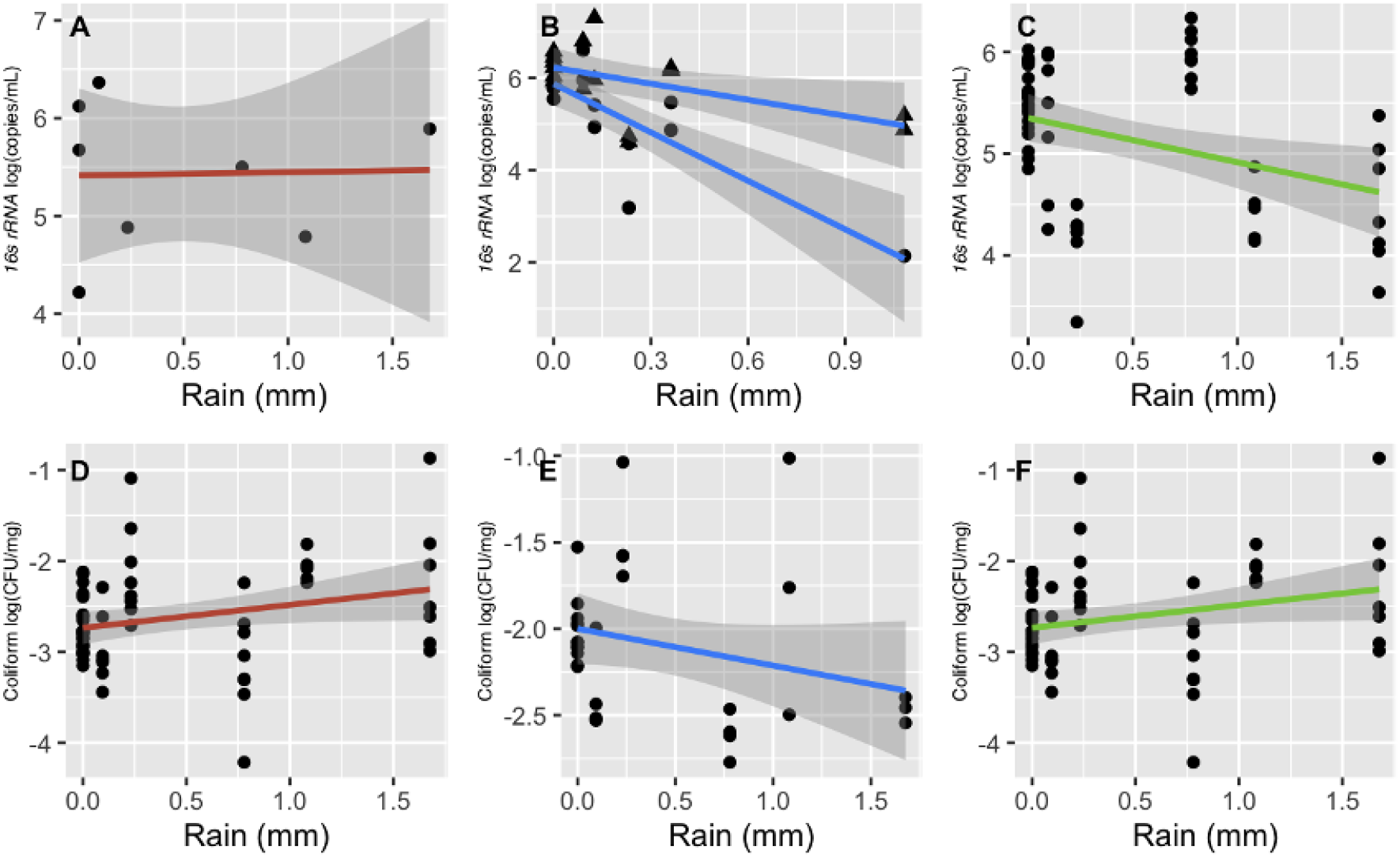
Effect of rain events on the concentration of sewage contaminants.

